# Protein-Protein Interaction Site Prediction Based on Attention Mechanism and Convolutional Neural Networks

**DOI:** 10.1101/2021.07.10.451856

**Authors:** Shuai Lu, Yuguang Li, Qiang Ma, Xiaofei Nan, Shoutao Zhang

## Abstract

Proteins usually perform their cellular functions by interacting with other proteins. Accurate identification of protein-protein interaction sites (PPIs) from sequence is import for designing new drugs and developing novel therapeutics. A lot of computational models for PPIs prediction have been developed because experimental methods are slow and expensive. Most models employ a sliding window approach in which local neighbors are concatenated to present a target residue. However, those neighbors are not been distinguished by pairwise information between a neighbor and the target. In this study, we propose a novel PPIs prediction model AttCNNPPISP, which combines attention mechanism and convolutional neural networks (CNNs). The attention mechanism dynamically captures the pairwise correlation of each neighbor-target pair within a sliding window, and therefore makes a better understanding of the local environment of target residue. And then, CNNs take the local representation as input to make prediction. Experiments are employed on several public benchmark datasets. Compared with the state-of-the-art models, AttCNNPPISP significantly improves the prediction performance. Also, the experimental results demonstrate that the attention mechanism is effective in terms of constructing comprehensive context information of target residue.

## 1 Introduction

PROTEIN-protein interactions play an import role in life processes by forming chemical bonds [1–4]. The bonding amino acid residues participating in interactions are protein-protein interaction sites (PPIs). Accurate recognition of these interaction sites is helpful for the development of novel therapeutics [5, 6], annotation of protein functions [7], and molecular mechanisms study of diseases [8, 9].

Judging a residue of protein belongs to the binding sites based on biological experimental methods remains costly and time-consuming [10–12]. In recent years, a lot of computational predictors for protein-protein interaction site (PPIs) prediction have been developed. These methods can be roughly divided into two groups: structure-based interface prediction and sequence-based interaction sites determination. Structure-based methods usually need the 3D structure of proteins [13–15]. Sequence-based methods only utilize information from protein sequence [16]. These methods usually solve the prediction problem in two ways: one is determining which residue pairs interact with each other from a set of given proteins and the other is judging which residues would interact with any other protein from a given protein sequence or structure [17]. Both types of methods rely on features extracted from sequence and/or structure such as evolutionary information [18], secondary structure [19], accessible surface area [20] and backbone flexibility [21]. A lot of computational techniques are used to utilize those features including shallow neural networks[20, 22, 23], support vector machine [24], random forest [25, 26], Bayesian techniques [27], ensemble learning [28] and deep learning [29–32].

Due to the rapid growth of unannotated protein sequence data, sequence-based approaches are becoming increasingly necessary [33]. In this study, we focus on the sequence-based methods and predict binding sites from a given protein sequence. A lot of sequence-based models employ a sliding window to gather information from the local environment of target residue [23, 27, 31, 34]. However, the sliding window approach doesn’t distinguish different neighboring residues for PPIs prediction by direct concatenation. In fact, different neighboring residue may play different roles. Inspired by a new direction of neural network research emerged in recent years [35], we employ attention mechanism to put different “attention” on different neighboring residues in the sliding window approach. The attention mechanism achieves success in machine translation [36], sentence classification [37], drug-target binding affinity prediction [38], paratope prediction [39] and protein structure prediction [40].

Here, we propose a novel deep learning model combing attention mechanism and convolutional neural networks (CNNs) for PPIs prediction. The attention mechanism is used to capture neighboring residue information and its correlation with the target residue. And, the CNNs are employed to extract information from local environment of target residue for prediction.

We name our method AttCNNPPISP short for **Att**ention-based **C**onvolutional **N**eural **N**etworks for **P**rotein-**P**rotein **I**nteraction **S**ites **P**rediction. The source code and datasets are available at https://github.com/biolushuai/attention-based-CNNs-for-PPIs-Prediction. The experiment is carried out on benchmark datasets, and AttCNNPPISP method out-performs the state-of-the-art methods. The detailed analyses show that attention mechanism is very important for PPIs prediction.

## 2 Materials and Methods

### 2.1 Datasets

In this study, we use the same protein sequences as the state-of-the-art method DeepPPISP [31] from three benchmark datasets Dset 186, Dset 72 [27] and PDBset 164 [34]. All the protein sequences in those datasets are collected form Protein Data Bank [41]. The sequences are derived from heterodimeric structures with resolution *<*3.0Å, sequence homology *<* 25% and length *>*50. Protein structures with missing ratio (i.e., the number of missing residues of a chain listed in REMARK465 divided by the total number of residues of the chain) *>*30% are removed. Also, protein complexes with interface area of *<*500Å^2^ or ≥2500Å^2^ as mentioned in PDBsum [42] and transmembrane proteins listed in PDBTM [43] are removed.

Same as works in previous studies [27, 31, 34], two conditions must be met if a residue is defined as an interaction site. First, it should be a surface residue and its relative solvent accessible surface area is less than 5%. Second, a surface residue is defined as an interaction site when it loses at least 1*Å*^2^ absolute solvent accessible surface area, before and after the binding form. Otherwise, the residue is defined as a non-interaction site.

These three datasets come from different resources, and are integrated to a fused dataset to ensure that training set and testing set are from an identical distribution, same as DeepPPISP [31]. The reconstruction of datasets is helpful for making full use of all proteins to train a deep learning model. Table 1 shows the size of datasets, the number of binding residues and non-binding residues.

**TABLE 1.** Summary of datasets.

| Datasets | NO. of Protein Sequences | NO. of Binding Residues | NO. of non-Binding Residues |
| --- | --- | --- | --- |
| TrainingSet | 352 | 11206 | 61982 |
| TestingSet | 70 | 2330 | 9461 |

### 2.2 Input Features

In this work, we use residue features including one-hot encoding of protein sequence, protein evolutionary information, and other predicted structural features from protein primary sequence. Because we focus only on the sequence-based computational method, the secondary structure information used here are predicted by NetsurfP-2.0 [44] rather than assigned by DSSP [45] which takes protein structure as input. We also take advantage of other features generated by NetsurfP-2.0, such as solvent accessibility and backbone dihedral angles. All those features are described in detail as follows:

#### 2.2.1 One-hot encoding of protein sequence

There are only 20 possible natural residue types, and we encode each raw protein sequence as a 20 dimensional one-hot vector, where each element is either 1 or 0 and 1 indicates the existence of a corresponding amino acid residue.

#### 2.2.2 Position-specific scoring matrix

Various related works have proved that the evolutionary information in position-specific scoring matrix (PSSM) is helpful for PPIs prediction [25, 28, 31]. By running PSI-BLAST [46] against the non-redundant (nr) [47] database with three iterations and an E-value threshold of 0.001, we get PSSM in which each residue is encoded as a 20D vector representing the probabilities of 20 natural amino acid residues occurring at each position. For each protein sequence with L residues, there are L rows and 20 columns in PSSM.

#### 2.2.3 Predicted structural features

In this study, we utilize NetsurfP-2.0 [44] to predict local structural features from protein primary sequence, including solvent accessibility, secondary structure, and backbone dihedral angles for each residue. NetsurfP-2.0 is a novel deep learning model trained on several independent datasets and achieves the state-of-the-art performance of predicting those local structural protein features.

For every reside in each input sequence, we calculate its absolute and relative solvent accessibility surface accessibility (ASA and RSA, respectively), 8-class secondary structure classification (SS8), and the backbone dihedral angles (*ϕ* and *ψ*). ASA and RSA represent the solvent accessibility of a residue. The predicted secondary structure describes the local structural environment of a residue. And, *ϕ* and *ψ* figure the relative positions of adjacent residues. The 8-class secondary structures are: 3-helix (G), a-helix (H), p-helix (I), b-strand (E), b-bridge (B), b-turn (T), bend (S) and loop or irregular (L). All those features are very helpful for PPIs prediction.

The state-of-the-art method DeepPPISP [31] utilizes a 49D feature vector including PSSM matrix, one-hot encoding of raw protein sequence and secondary structure calculated by DSSP [45]. However, predicted structural features consisting of solvent accessibility, secondary structure, and backbone dihedral angles are used in our work, and each residue in every input protein sequence is represented as a 52D feature vector. Features used in this study are richer than those in DeepPPISP.

### 2.3 Model architecture

The PPIs prediction problem can be summarized as a binary classification task: judging whether a residue from a given protein sequence binding with its partner protein or not. As described in Section 2.2, each residue is encoded into a 52D vector. And each protein sequence can be represented as a matrix S, including a list of residues:

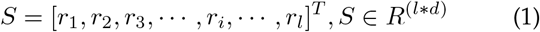

where *r*_*i*_ ∈ *R*^*d*^ is the residue feature vector corresponding to the *i*-th residue in the protein sequence, *l* is the protein sequence length, and *d* is the residue feature dimension (52 in this paper).

#### 2.3.1 Convolution Neural Networks

Convolution Neural Networks (CNNs) model has been adapted to various bioinformatics tasks such as protein binding site prediction [48], protein-ligand scoring [49] and protein-compound affinity prediction [50]. For extracting features from residue’s local environment information, we utilize a CNNs model for PPIs prediction. CNNs model in this study has similar structure as TextCNN [51] in DeepPPISP [31] but small convolution kernel sizes. And, to explore the effects of attention mechanism, AttCNNPPISP utilizes the same CNNs for comparison.

In this study, the input of CNNs is a protein sequence represented as a matrix *S*. The CNNs model employs a convolutional operation on a sliding window of length *w*(*w* = 2*n* + 1), where *n* could be any positive integer for the *i*-th target residue *r*_*i*_. It means that we consider the target amino acid at the center and 2*n* neighboring residues as input features of the target residue. Similar as DeepPPISP, we use the all-zero vector padding for those residues which do not have neighboring residues in the left or right. A convolutional operation can be shown as:

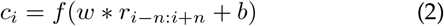

where *f* is a non-linear activation function, and *r*_*i−n*:*i*+*n*_ denotes the concatenation of *w* residue vectors:*r*_*i−n*:*i*+*n*_ = [*r*_*i−n*_, *r*_*i−n*+1_, …, *r*_*i*_, …, *r*_*i*+*n−*1_, *r*_*i*+*n*_], where *r*_*i*_ is the target residue, and *r*_*i−n*_, *r*_*i−n*+1_, …, *r*_*i−*1_, *r*_*i*+1_, …, *r*_*i*+*n*_ are neighboring residues. After convolutional operation, a max pooling operation is applied on *c*_*i*_, and two fully connected layers are followed which predicts the interaction probability of the target residue. However, CNNs model treats neighboring residues within a sliding window equally by direct concatenation, and ignores their different effects on the target residue.

#### 2.3.2 Attention-based Convolution Neural Networks

As shown in Fig.1, AttCNNPPISP consists of four parts, i.e., input layer, attention layer, convolution & pooling layer and fully-connected layer. The input layer takes the protein sequence as input and extract all sequence-based features. The convolution & pooling and fully-connected layers share the same architecture as CNNs model said in Section 2.3.1. The attention layer utilizes attention mechanism which is employed as an additional fully-connected layer, and it is trained in conjunction with all the other network components like other machine learning works [37]. To be more specific, the attention layer is to build a context vector representation for each residue. The context vector is concatenated with the input residue vector to form a new residue representation, and it is then fed to the convolution & pooling layer.

**Fig. 1.**
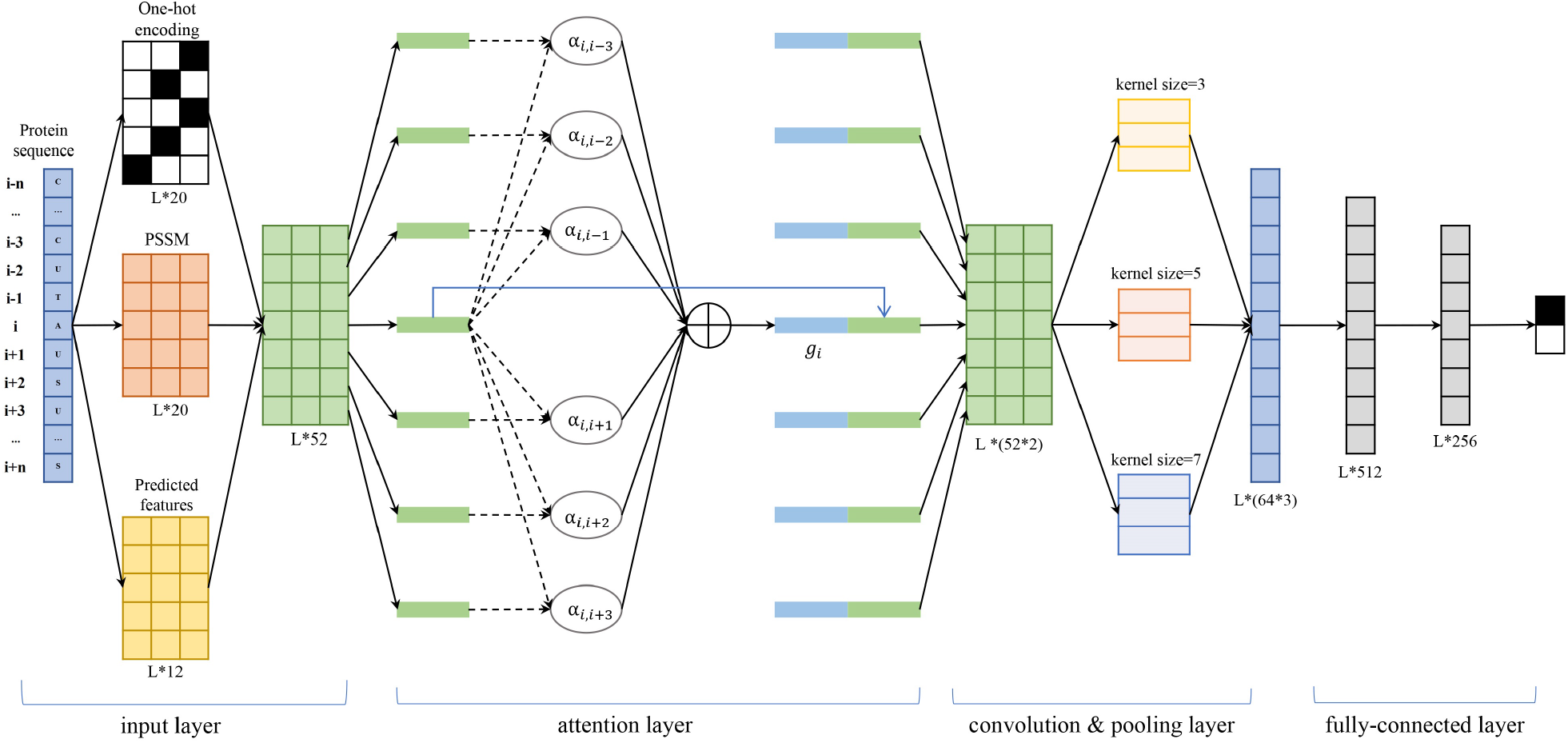
The deep neural network architecture of AttCNNPPISP. In attention layer, the dotted lines correspond to formulas (3)-(4), the black solid lines correspond to formula (5), the blue solid line is copy operation. The input and output dimensionality of each layer and kernel sizes in convolution & pooling layer are marked.

The attention mechanism enables a target residue learning to pay different attention on each neighboring residue. Here we utilize typical additive model. And, the similarity score *score*(*r*_*i*_, *r*_*j*_) between target residue *r*_*i*_ and neighboring residue *r*_*i*_ and attention weights *α*_*i,j*_ are calculated as:

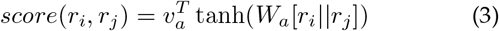

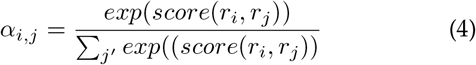

The residue pair correlation score *score*(*r*_*i*_, *r*_*j*_) is computed by a two-layer neural network described in formula (3). And neighboring residues *r*_*j,j≠i*_ with larger scores contribute more on context vector *g*_*i*_. The attention weights *α*_*i,j*_ should satisfy the following restrictions: *α*_*i,j*_ ≥ 0 and Σ_*i*_ *α*_*i,j*_ = 1.

For constructing the context vector *g*_*i*_, neighboring residue vectors are scored and combined in a weighted sum:

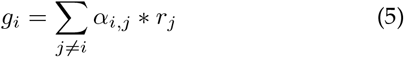

After attention layer, each residue *r*_*i*_ is concatenated with its context vector *g*_*i*_ to form an extended residue vector 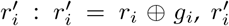 is then used to fed into the convolution & pooling layer. Attention mechanism determines which residues should be put more attention on than other residues over the sequence in PPIs prediction. At last, the fully-connected layer output the binding probability of each target residue.

### 2.4 Performance assessment

We use six evaluation metrics to evaluate the performances of the PPIs prediction models. Five of them are classic threshold-dependent binary classification metrics as shown in the following:

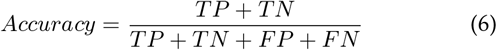

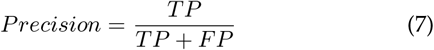

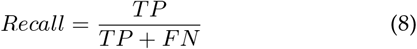

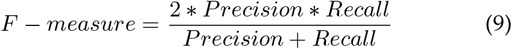

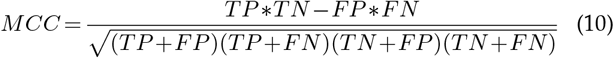

where, TP (True Positive) is the number of interacting residues that are correctly predicted as interacting, FP (False Positive) is the number of non-interacting residues that are falsely predicted as interacting, TN (True Negative) denotes the number of non-interacting sites that are identified correctly, and FN (False Negative) denotes the number of interacting sites that are identified falsely. In this work, all those five metrics are calculated using a threshold for maximizing F-measure [31].

Because all those five metrics are threshold-dependent, we also utilize the area under the precision-recall curve (AUPRC) which gives a threshold-independent evaluation on the overall performance. Also, AUPRC is sensitive on imbalanced data, and PPIs prediction is normally an imbalanced learning problem [31]. Therefore, we take AUPRC as the most import metric for model evaluation [52].

### 2.5 Implementation

We implement AttCNNPPISP using PyTorch. The training configurations are: loss function: cross-entropy loss, optimization: Adaptive Momentum (Adam); learning rate: 0.1, 0.01, 0.001; batch size: 32, 64, 128; dropout: 0.2, 0.5, 0.7; sliding window length: 3, 5, 7, 9, 11. An independent validation set (10% of our training set) is used to tune parameters. For each combination, networks are trained until the performance on the validation set stops improving or for a maximum of 150 epochs. Training time of each epoch varies roughly from 5 to10 minutes depending on the sliding window length, using a single NVIDIA RTX2080 GPU.

The input channel and out channel of convolution & pooling layer are 1 and 64, respectively. Three kernel sizes (3, 5 and 7) are used, and the results are concatenated and fed to a two-layer fully connected network. The first fully connected layer has 512 nodes and the second fully connected layer has 256 nodes.

## 3 Results

### 3.1 Comparison with competing methods

To evaluate the performance of AttCNNPPISP for PPIs prediction, we compared it with six competing computational methods including PSIVER [27], SPPIDER [20], SPRINGS [34], ISIS [23], RF PPI [25] and DeepPPISP [31]. It must be noted that DeepPPISP [31] uses not only protein sequence but also structure information when it computes protein secondary structure using DSSP [45] which takes protein structure as input.

Except for DeepPPISP utilizing a deep learning framework, all others use shallow machine learning methods. The six competing methods all employ a sliding window approach. The sequenced-based features used in those models include PSSM and predicted properties such as solvent accessibility, fingerprints, evolutionary information, and structural information. AttCNNPPISP uses one-hot encoding of protein sequence, PSSM, and predicted accessibility, secondary structure information and backbone dihedral angles. Specially, DeepPPISP combines local features with global feature from the whole protein sequence. AttCNNPPISP only utilizes local features from residues within a sliding window length.

Table2 shows the results of AttCNNPPISP and six competing methods on the testing set. Results of PSIVER, SPPIDER, SPRINGS, ISIS, RF PPI and DeepPPISP are excerpted from DeepPPISP [33]. For AttCNNPPISP, we use a threshold of 0.21 determined as explained in Section 2.4. AttCNNPPISP outperforms other competing methods on most assessment metrics. Although the ACC of AttCNNPPISP is slightly lower than ISIS, it achieves the highest scores on all other metrics. Especially for AUPRC, AttCNNPPISP apparently obtains higher result than all five competing methods and improves by 3.9 percentage points over the state-of-the-art method DeepPPISP.

**TABLE 2.** Performance of AttCNNPPISP and other competing methods.

| Method | AUPRC | ACC | Precision | Recall | F-measure | MCC |
| --- | --- | --- | --- | --- | --- | --- |
| PSIVER | 0.250 | 0.653 | 0.253 | 0.468 | 0.328 | 0.138 |
| SPPIDER | 0.230 | 0.622 | 0.209 | 0.459 | 0.287 | 0.089 |
| SPRINGS | 0.280 | 0.631 | 0.248 | 0.598 | 0.350 | 0.181 |
| ISIS | 0.240 | **0.694** | 0.211 | 0.362 | 0.267 | 0.097 |
| RF_PPI | 0.210 | 0.598 | 0.173 | 0.512 | 0.258 | 0.118 |
| DeepPPISP | 0.320 | 0.655 | 0.303 | 0.577 | 0.397 | 0.206 |
| AttCNNPPISP | **0.359** | 0.657 | **0.313** | **0.611** | **0.414** | **0.229** |

Fig2 visually display the true and predicted interaction sites by DeepPPISP and AttCNNPPISP on a representative protein (PDB ID: 1JTD, Chain: B). This protein is consisting of 273 residues and has 33 interaction sites. The webserver of DeepPPISP at http://bioinformatics.csu.edu.cn/PPISP/ is not accessible. To make a fair comparison, we download the source code of DeepPPISP from https://github.com/CSU-BioGroup/DeepPPISP and run it on the same local hardware and software environment with AttCNNPPISP. From Fig2, we can observe that AttCNNPPISP predict more true positive and less false positive interaction sites.

**Fig. 2.**
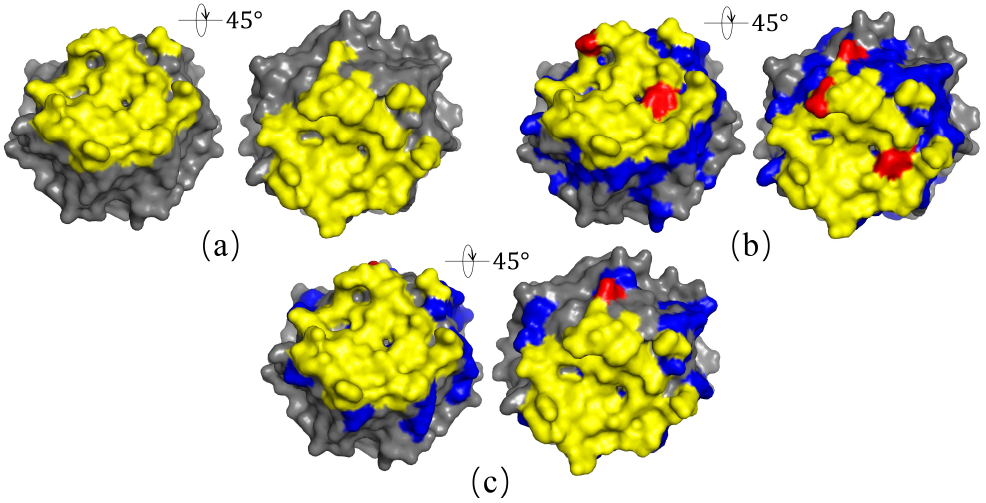
Interaction sites of a representative protein in the testing set (PDB ID: 1JTD, Chain: B). (a) True interaction sites are in yellow on the protein surface which are labeled in the testing set. (b) Interaction sites predicted by DeepPPISP. (c) Interaction sites predicted by AttCNNPPISP. In figure b and c, TP predictions are in yellow, FN predictions are in red, FP predictions are in blue and the background grey represents TN predictions.The left and right images of figures b and c show two different viewing angles (i.e., 45° rotated view).

### 3.2 The effects of different sliding window lengths

We employ the sliding window approach with different lengths (i.e., 3, 5, 7, 9, 11). As shown in Table 3, AUPRC, Precision, Recall, F-measure and MCC obtained by AttCNNPPISP of a sliding window length 5 are 0.359, 0.657, 0.313, 0.414 and 0.229, respectively, which are better than all other window lengths. Recall of window length 5 is lower than window lengths 3, 7 and 11. However, Precision and Recall are usually contradictory performance metrics. The higher Precision is always accompanied by the lower Recall. Therefore, the results show that AttCNNPPISP performs best when the sliding window length is 5.

**TABLE 3.**
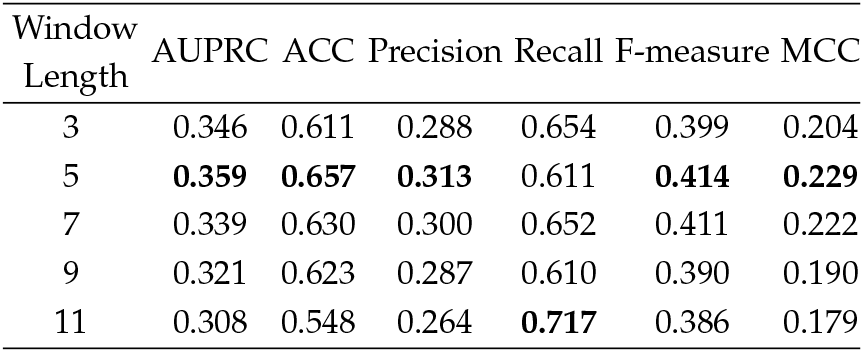
Performance of AttCNNPPISP with different sliding window length.

| Window Length | AUPRC | ACC | Precision | Recall | F-measure | MCC |
| --- | --- | --- | --- | --- | --- | --- |
| 3 | 0.346 | 0.611 | 0.288 | 0.654 | 0.399 | 0.204 |
| 5 | **0.359** | **0.657** | **0.313** | 0.611 | **0.414** | **0.229** |
| 7 | 0.339 | 0.630 | 0.300 | 0.652 | 0.411 | 0.222 |
| 9 | 0.321 | 0.623 | 0.287 | 0.610 | 0.390 | 0.190 |
| 11 | 0.308 | 0.548 | 0.264 | **0.717** | 0.386 | 0.179 |

For determining a suitable window size, PISVER uses lengths from 3 to 21 and achieves the highest performance when the window length is 9 And, DeepPPISP applies lengths from 7 to 15 and obtains the best results when the window length is 7. The results of PISVER and DeepPPISP as well as Table 3 shows that too long or too short sliding window length isn’t helpful for PPIs prediction, and an appropriate number of neighboring residues is effective for better representation of target residue’s local environment. When the sliding window length is 3, the performance of AttCNNPPISP is poor as there are too few neighboring residues to get effective information of target residue for prediction. When the sliding window length is too long, such as 11, too many neighboring residues cause distraction of attention.

### 3.3 The effects of different feature types

To find out what role each feature type plays in AttCNNPPISP, we compare the performances of AttCNNPPISP with a sliding window length of 5 using different feature combinations. Table 4 shows the effects of different input feature types in AttCNNPPISP. AUPRC drops to 0.347, 0.303, 0.353, 0.279, and 0.339 from 0.359, when the input features of AttCNNPPISP miss One-hot, PSSM, SS8, ASA and RSA, and backbone dihedral angles (*ϕ* and *ψ*), respectively. Among them, input features missing ASA and RSA have the largest drop. When the missing feature is ASA and RSA, ACC, Precision, Recall, F-measure, MCC drop to 0.544, 0.248, 0.641, 0.358 and 0.129, respectively. Except for ACC, the missing of ASA and RSA cause largest drop on other metrics. It indicates that ASA and RSA are the most import features in AttCNNPPISP.

**TABLE 4.**
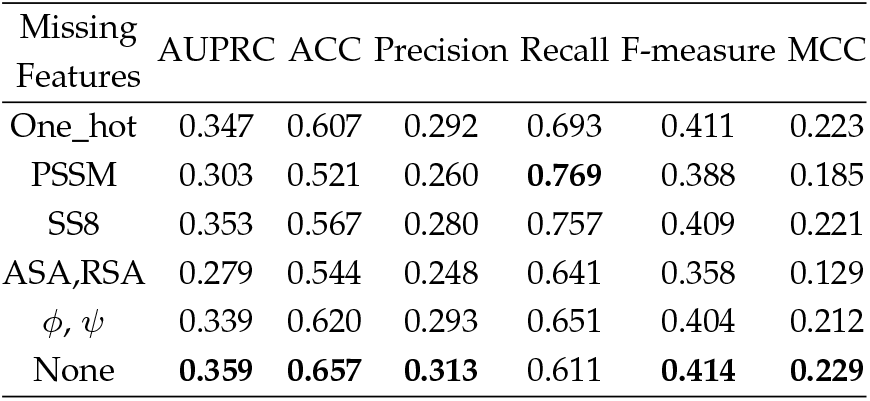
Performance of AttCNNPPISP with different features combination.

According to DeepPPISP [31], the raw protein sequence, PSSM, secondary structure are used and raw protein sequence contributes most. Different identification of the most important feature is caused by the different model architectures. In our study, the usage of new predicted features provides more useful information and get more attention. And the same conclusion is that all features used can get best results.

### 3.4 The effect of attention mechanism

As said in Section 2.3.1, we use a CNNs model for comparison to explore the effects of attention mechanism. We apply the CNNs model on the same datasets and Table 5 shows that AttCNNPPISP significantly outperforms the CNNs model on most evaluation metrics due to the attention mechanism.

**TABLE 5.** Effect of attention mechanism.

| Method | AUC PR | ACC | Precision | Recall | F-measure | MCC |
| --- | --- | --- | --- | --- | --- | --- |
| CNNs | 0.350 | 0.641 | 0.304 | **0.631** | 0.410 | 0.222 |
| AttCNNPPISP | **0.359** | **0.657** | **0.313** | 0.611 | **0.414** | **0.229** |

Fig3 shows the attention score using color gradient between a target residue 220 with its four neighboring residues of a representative protein. This residue is an interaction site rightly predicted by AttCNNPPISP. The target residue 220 and its neighboring residues are all correctly predicted. Interaction sites are shown in green and non-interaction sites are shown in black.

**Fig. 3.**
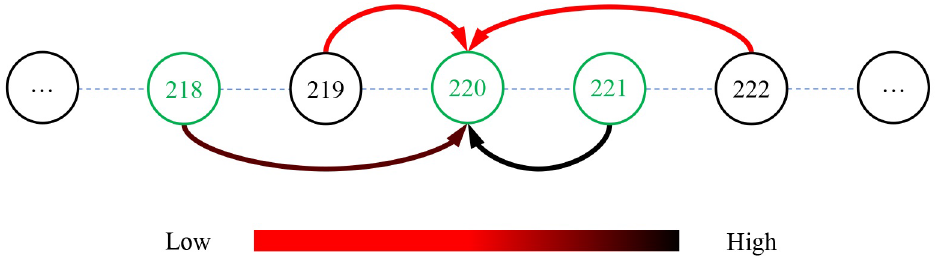
Attention scores generated by AttCNNPPISP with sliding window length 5 for a representative residue of a protein (PDB ID: 1JTD, Chain: B, Target residue: 220). The neighboring residues in green are the interaction sites labeled in the dataset.

From Fig3, we can see that the attention scores between interaction sites are high (darker color) which is consistent with biological views and has useful information. As expected, the attention mechanism distinguishes different effect of each neighboring residue and provides more accurate information about target residue’s local environment.

Fig4 shows the averaged attention score of each residue type pair with various sliding window sizes. The vertical axis represents the 20 types of target residues and the horizontal axis represents the neighboring residues. All residue types are in alphabetical order. The red color represents that more attention is put on the corresponding neighboring residue. On the contrary, the green color represents that less attention is put. For example, when the sliding window length is 11, and the target residue is Cysteine (C), the residue Tryptophan (W) draws the most attention among all neighboring residue types.

**Fig. 4.**
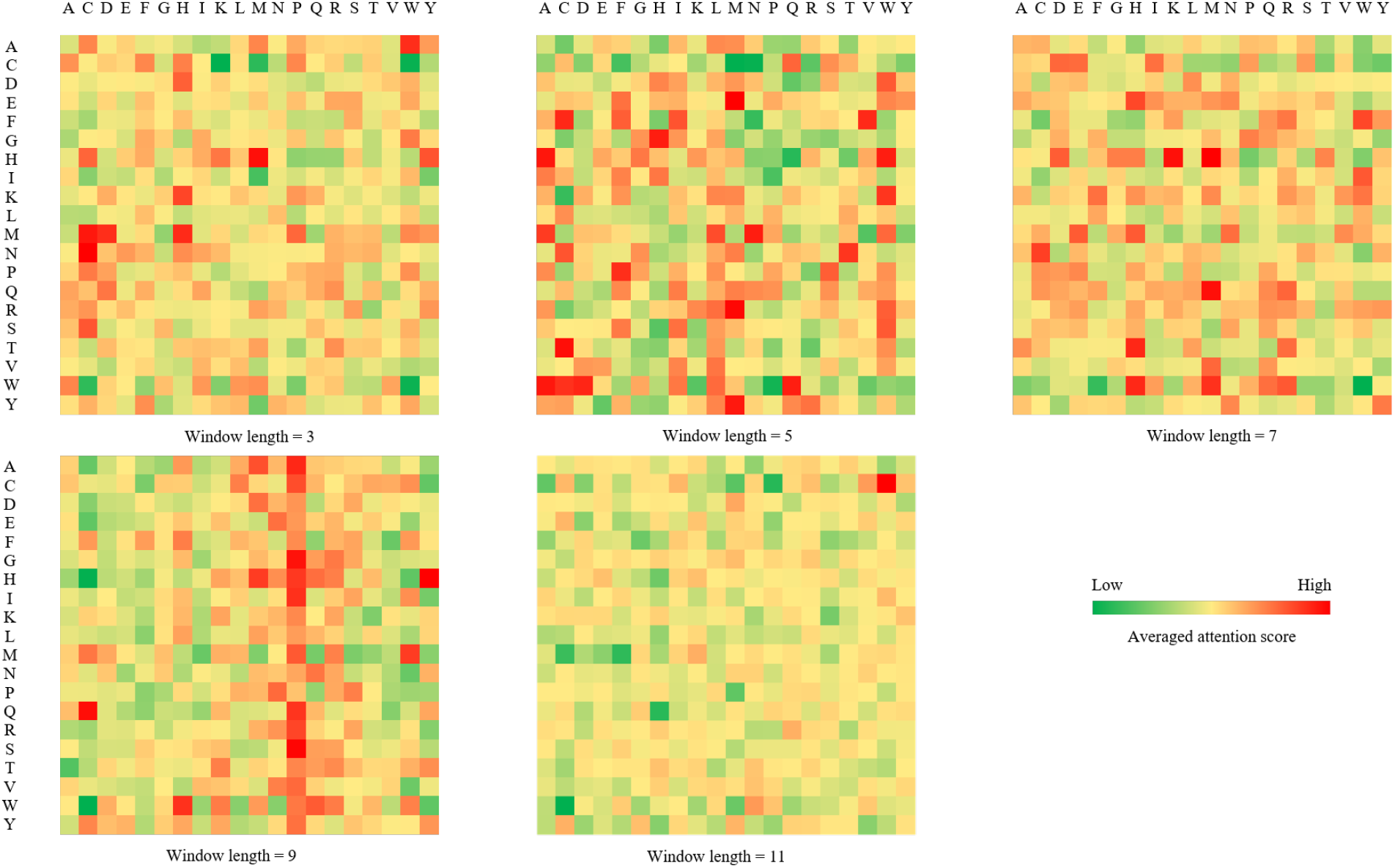
Attention visualization of each residue type pair with various sliding window lengths.

The results show that the variations in attention scores don’t correlate with the properties of amino acid residues such as polarity, charge and alkalinity or acidity. That is, the positively charged residue does not always attract the attention of the negatively charged residue. The same goes for other properties of residues. The reason it that the attention mechanism captures context information of protein sequence other than properties of individual residue in our model.

However, it is obvious that when the sliding window length is 3, there are very few dark red and dark green cells. In other words, neighboring residues don’t get specific and effective attention as the window length is too short. From both Table3 and Fig4, we can see that when the sliding window length is 5, our model performs best and neighboring residues get most specific and effective attention. And when the length of the sliding window grows longer than 5, the averaged attention scores become even. In particular, when the sliding window length is 11, almost every neighboring residue doesn’t get enough attention. When the sliding window length is 9, attention focuses on a few types of neighboring residues such as Proline (P) which cause poor results. We count the number of occurrences of each residue type which is not relevant to this concentration of attention. This situation needs further study.

### 3.5 Running time

We investigate the time required to generate the features used by DeepPPISP and AttCNNPPISP and the time required for prediction. All analyses were run on the same machine with Intel core i9-9900X CPU (10 cores), 64GB RAM, NVIDIA GeForce RTX 2080 GPU (8GB memory).

AttCNNPPISP is a bit lower than DeepPPISP. Table 6 shows feature generation time on the shortest (51 residues) and the longest (376 residues) protein sequences in the testing set. Both methods utilize PSSM returned by PSI-BLAST [46] which is time-consuming. As we focus on the sequenced-based approach and use structural features predicted by NetsurfP-2.0 [44], AttCNNPPISP takes about 300 seconds more to process a protein sequence of length 375. And DeepPPISP consists of a relatively simple neural network structure, it takes less time to train. Most of all, more time is spent in feature generation and training model, but AttCNNPPISP and DeepPPISP take similar time to make predictions.

**TABLE 6.** Running time comparison between DeepPPISP and AttCNNPPISP

| Protein Length | Method | Feature geneation | Inference |
| --- | --- | --- | --- |
| 376 | DeepPPISP | 4494.19 | 13.63 |
|  | AttCNNPPISP | 4810.07 | 14.15 |
| 51 | DeepPPISP | 3028.73 | 11.02 |
|  | AttCNNPPISP | 3170.16 | 11.84 |

## 4 Conclusion

In this study, we propose a deep learning model named AttCNNPPISP for sequence-based PPIs prediction. AttCNNPPISP combines attention mechanism with convolutional neural networks to capture the residue correlations within a sliding window. AttCNNPPISP can distinguish the effects of different neighboring residues on a target residue, and it captures accurate local environment information of the target residue. The experiments show that AttCNNPPISP significantly outperforms other state-of-the-art methods.

Though AttCNNPPISP has superior performance over other competing methods, it also has some potential limitations. First, similar like other sequence-base methods, our program takes a lot of time to generate sequence profiles by running PSI-BLSAT [46] and NetsurfP-2.0 [44]. Second, although the attention mechanism improves the overall performance, it takes additional computation consumption to calculate attention scores, especially when the sliding window length is long. Third, with the sliding window length growing, the attention mechanism loses focus and doesn’t lead to better results.

In this study, we show that paying different attention on different neighboring residue can be helpful for sequence-based PPIs prediction. We believe that the attention mechanism has a great potential in other biological sequence analysis and prediction problems. Currently we only employ one attention layer, and more attention layers may work better if sufficient data samples are available.

However, amino acid residues that are far apart in the sequence may be adjacent in space, and longer sliding window length needs to be considered. The experimental results show than too big sliding window may cause the distraction of attention. This problem needs further study and discussion.

## Acknowledgments

This work was supported by Bingtuan Science and Technology Project(2019AB034), ‘Created Major New Drugs’ of Major National Science and Technology (No. 2019ZX09301-159), Leading Talents Fund in Science and Technology Innovation in Henan Province(194200510002), and Natural Science Foundation of Henan Province of China (Grant No.202300410381). Xiaofei Nan and Shoutao Zhang are the corresponding authors for this paper.

---

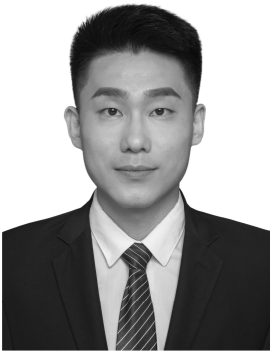

**Shuai Lu** received the BS degree from the School of Information Engineering of Zhengzhou University, Zhengzhou, Henan, China in 2014. Currently, he is working towards the PhD degree in the School of Computer and Artificial Intelligence of Zhengzhou University. His current research interests mainly include machine learning and data mining applied to the biological field, especially in the prediction of protein-protein and antibody-antigen interactions.

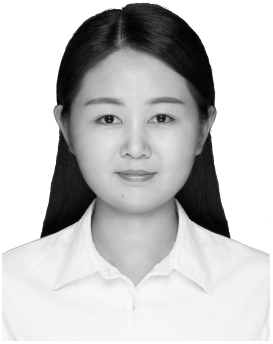

**Yuguang Li** received the MS degree from the School of Information Engineering, Zhengzhou University, Zhengzhou, Henan, China in 2018. Her research interests include bioinformatics, data mining, and NLP.

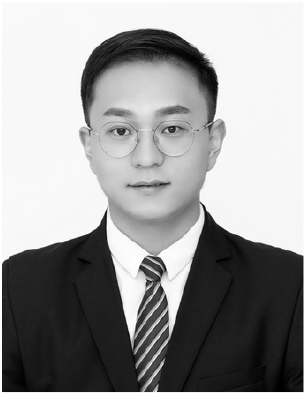

**Qiang Ma** received the PhD degree from Wuhan University. He has published 12 high-quality papers in journals. He is a lecturer in college of Life Sciences, Zhengzhou University, Henan province. His scientific interests are mainly in molecular virology, including bioinformatics.

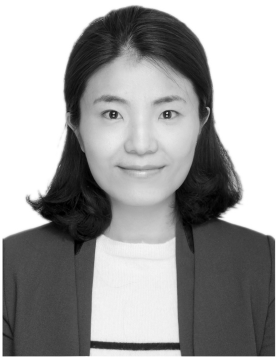

**Xiaofei Nan** received the PhD degree from the University of Mississippi. She is an associate professor in the School of Computer and Artificial Intelligence, Zhengzhou University, Henan. Her current interests include pattern recognition, data mining, and bioinformatics.

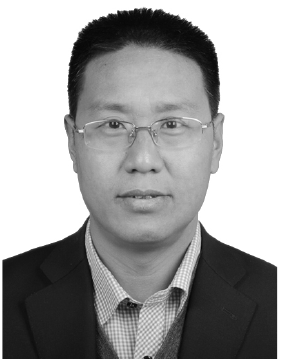

**Shoutao Zhang** received the PhD degree from the Northwest A&F University. He has published about 40 high-quality referred papers in international conferences and journals. He is a professor in the School of Life Sciences, Zhengzhou University, Henan. His research interests include bioinformatics.

